# New tools to monitor *Pseudomonas aeruginosa* infection and biofilms *in vivo* in *C. elegans*

**DOI:** 10.1101/2024.08.09.607303

**Authors:** Martina Ragno, Feng Xue, Sarah A. Blackburn, Michael Fasseas, Sushmita Maitra, Frederique Tholozan, Rachel Thompson, Laura Sellars, Rebecca Hall, Chris Saunter, David Weinkove, Marina Ezcurra

**Affiliations:** School of Biosciences, University of Kent, Canterbury, CT2 7NJ, United Kingdom; Magnitude Biosciences Limited, NETPark Plexus, Thomas Wright Way, Sedgefield, TS21 3FD, UK; Perfectus Biomed Group, Sci-Tech Daresbury, Keckwick Lane, Chesire, WA4 4AD, UK; Department of Biosciences, Durham University, Stockton Road, Durham, DH1 3LE, UK

**Keywords:** Pseudomonas aeruginosa, biofilms, quorum sensing, C. elegans, antimicrobial resistance

## Abstract

Antimicrobial resistance is a growing health problem. *Pseudomonas aeruginosa* is a pathogen of major concern because of its multidrug resistance and global threat, especially in health-care settings. The pathogenesis and drug resistance of *P. aeruginosa* depends on its ability to form biofilms, making infections chronic and untreatable as the biofilm protects against antibiotics and host immunity. A major barrier to developing new antimicrobials is the lack of *in vivo* biofilm models. Standard microbiological testing is usually performed *in vitro* using planktonic bacteria, without representation of biofilms, reducing translatability. Here we develop tools to study both infection and biofilm formation by *P. aeruginosa in vivo* to accelerate development of strategies targeting infection and pathogenic biofilms. Using the nematode *Caenorhabditis elegans* and *P. aeruginosa* reporters combined with *in vivo* imaging we show that fluorescent *P. aeruginosa* reporters that form biofilms *in vitro* can be used to visualise tissue infection. Using automated tracking of *C. elegans* movement, we find that that the timing of this infection corresponds with a decline in health endpoints. In a mutant strain of *P. aeruginosa* lacking RhlR, a transcription factor that controls quorum sensing and biofilm formation, we find reduced capacity of *P. aeruginosa* to form biofilms, invade host tissues and negatively impact healthspan and survival. Our findings suggest that RhlR could be a new antimicrobial target to reduce *P. aeruginosa* biofilms and virulence *in vivo* and *C. elegans* could be used to more effectively screen for new drugs to combat antimicrobial resistance.

## BACKGROUND

The Gram-negative bacterium *Pseudomonas aeruginosa*, a common and opportunistic pathogen, causes disease in a variety of hosts(Rahme et al., 1997; S et al., 2011; Tan et al., 1999). In humans, *P. aeruginosa* can cause serious complications, form systemic infections in immunodeficient patients and develop into chronic infections(Govan and Deretic, 1996; Lieberman and Lieberman, 2003; Lyczak et al., 2000). *P. aeruginosa* infections are characterised by antibiotic resistance, limited treatment options and high mortality, and outbreaks caused by multidrug resistant strains are on the rise (Fisher et al., 2005; Obritsch et al., 2005; R et al., 2001). Pathogenesis and drug resistance of *P. aeruginosa* depends on its ability to form biofilms, which make infections chronic and untreatable as the biofilm protects against antibiotics and host immunity(Roy et al., 2018). Although biofilms are central to chronic *P. aeruginosa* infections, research examining *P. aeruginosa* virulence has largely focused on planktonic bacteria or biofilm studies performed *in vitro* or *ex vivo*, reducing translatability and creating a barrier to the development of effective antimicrobials to limit chronic *P. aeruginosa* infections(Highmore et al., 2022).

Biofilm-related infections are challenging to study and monitor *in vivo* in because they are typically internal. While e.g. surface wounds can be monitored in real time, analysis of internal biofilms are typically carried out postmortem on *ex vivo* tissue. There are a few examples of *in situ in vivo* monitoring of biofilms using advanced imaging techniques, such as microcomputed tomography and micropositron emission tomography but these techniques are highly specialised and expensive(Guzmán-Soto et al., 2021). Developing novel *in vivo* approaches that allow high-throughput assays to study biofilm-related infections would increase the understanding of bacterial biofilms and interactions between biofilms and host, and could lead to the development of new strategies targeting antimicrobial resistance and biofilms.

The model organism *Caenorhabditis elegans* offers a valuable tool to study infection that can be developed to perform detailed studies of biofilm formation *in vivo*, increasing the understanding of bacterial physiology within biofilms and interactions between biofilms and the host. *C. elegans* has a small size and rapid generation time, can be used to conduct large-scale screening and offers an extensive research toolkit including functional assays to monitor health outputs and physiology. *C. elegans* is susceptible to human pathogens and *C. elegans-P. aeruginosa* infection models are particularly useful, as many *P. aeruginosa* virulence-related factors are conserved across widely divergent taxa from nematodes to plants to mammals(Kim and Ausubel, 2005; Rahme et al., 1995; Tan and Ausubel, 2000), and the human innate immune system shares many characteristics with that of *C. elegans*. In addition, *C. elegans* is transparent, making it possible to image fluorescent reporters and infection in living animals in real time. Thus, *C. elegans* has potential to be developed as a new model to study pathogenic biofilms *in vitro*. Here we report that quorum sensing (QS), a signalling process by which *P. aeruginosa* regulates biofilm formation, is required for *P. aeruginosa* pathogenicity in *C. elegans*.

QS enables bacteria to respond to changes in the density and composition of the surrounding bacterial community and synchronise behavioural responses through extracellular molecules, autoinducers, to produce biofilms(Thi et al., 2020). In *P. aeruginosa*, QS is regulated through the production and secretion of the autoinducers *N*-3-oxo-dodecanoyl-L-homoserine lactone (3O-C_12_-HSL) and *N*-butyryl-L-homoserine lactone (C_4_-HSL), which are produced by the canonical acylated homoserine lactone synthases LasI and RhlI, respectively. 3O-C_12_-HSL is sensed by the transcriptional regulator LasR and C_4_-HSL is sensed by the transcriptional regulator RhlR. Binding of the autoinducers to LasR and RhlR results in transcriptional responses and expression of genes important for biofilm formation and virulence(Mukherjee et al., 2017; O’Loughlin et al., 2013; Thi et al., 2020) (Figure 1a).

**Figure 1.**
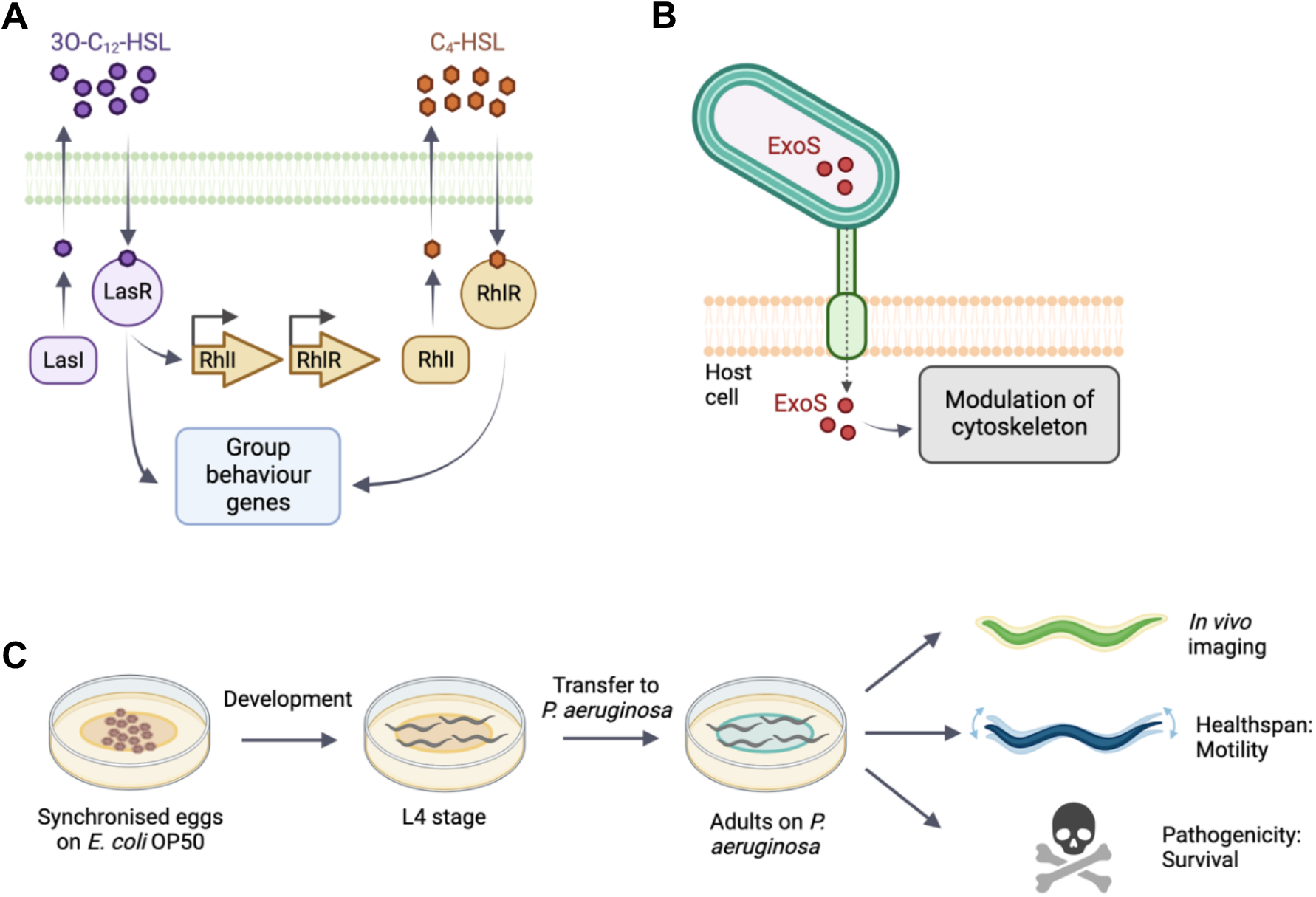
Schematic of quorum sensing and type-III secretion in *P. aeruginosa* and *C. elegans* experimental setup. **a**) The acyl-homoserine lactone-based quorum sensing network. LasI produces and LasR responds to the autoinducer 3OC12-HSL. The LasR:3OC12–HSL complex activates transcription of many genes including rhlR. RhlI generates the autoinducer C4-HSL. **b**) *P. aeruginosa* injects exotoxin ExoS though a needle complex after contact with the surface of the targeted eukaryotic cell. ExoS modulates the cytoskeleton. **c**) *C. elegans* populations are synchronised and cultivated on *E. coli* OP50. At L4 stage, animals are transferred onto the different experimental bacterial strains. Bacterial fluorescence, healthspan and survival are monitored throughout adulthood.

In this study, we use fluorescent reporters for the RhlI/R QS system to examine and monitor infection, and the resulting pathogenicity *in vivo* in *C. elegans*. We also examine a reporter of the virulence factor exoenzyme S (ExoS), a type III secretion effector which is injected into the host cell during infection, where it reorganises the cytoskeleton and induces apoptosis (Figure 1b). The type III secretion system is important for acute phases of infection but a role in biofilm formation has not been clearly established(Jouault et al., 2022).

Using fluorescent *P. aeruginosa* reporters, we confirm that QS is required for *P. aeruginosa* to form biofilms *in vitro*. We combine *in vivo* epifluorescence microscopy and functional readouts using automated tracking systems (Figure 1c) to demonstrate that RhlR signalling is required for *P. aeruginosa* to invade *C. elegans* tissues, leading to a reduction in healthspan and survival. Our findings suggest that *P. aeruginosa* infects *C. elegans* through QS and biofilm formation, and that methods combining *C. elegans*, functional assays and bacterial fluorescent reporters can be developed into high-throughput approaches to investigate pathogenic biofilms and evaluate anti-biofilms strategies *in vivo*.

## METHODS

### Bacterial strains and growth conditions

Bacterial strains used in this study are listed in Table 1. Bacterial strains were grown in Luria-Bertani broth (LB) and on LB plates fortified with 1.5% Bacto agar at 37°C. Single bacterial colonies were inoculated and incubated at 37°C for 16 hours at 250rpm. When appropriate, antimicrobials were included at the following concentrations: 200 μg/mL carbenicillin, 25 μg/mL gentamycin, 50 μg/mL tetracycline. For assays using 4-Nitropyridine-N-oxide (NPO), the compound was dissolved in DMSO and added to bacterial cultures, keeping DMSO at a maximum of 1% (v/v).

**Table 1.**
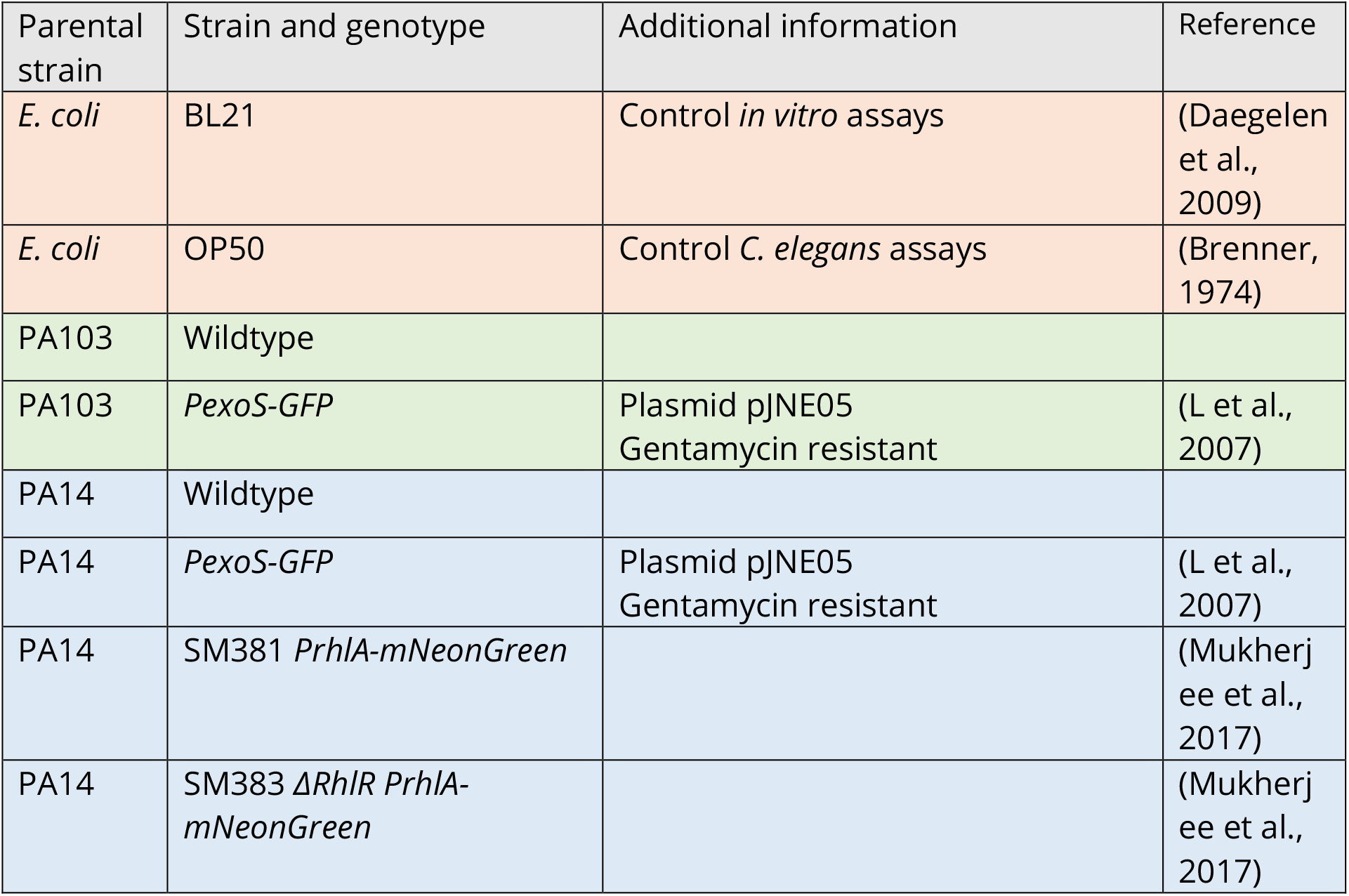
Bacterial strains used in study.

### Crystal violet staining assay

Static biofilm formation was evaluated using the crystal violet staining method as previously described(Campo-Pérez et al., 2023). Briefly, bacterial cultures were washed with PBS three times to remove antibiotics, resuspended in Mueller Hinton broth (M-HB) and diluted to OD600 0.2. 10 μl of each sample was mixed with 190 μl of M-HB in 96-well microplate. After 2 hours of static incubation at 37°C media and non-adhered cells were removed and replaced with fresh M-HB and further incubated for 24 hrs. Adherent cells were quantified by staining with crystal violet and measurement of A_550_. The assay was performed with three replicates for each condition.

### Fluorescence biofilm assay

96-well microplates with bacterial cultures were prepared as for the Crystal Violet assay but instead of staining GFP fluorescence was measured using a microplate spectrofluorometer reader. The assay was performed with three replicates for each condition.

### *C. elegans* culture and strains

*C. elegans* were maintained at 15°C on media plates seeded with *E. coli* OP50. For epifluorescence imaging and survival experiments, animals were cultivated using plates with 10 mL Nematode Growth Medium. For healthspan assays, plates with 15 mL defined agar were used(Maynard et al., 2018). The *C. elegans* strains used in this study was SS104 *glp-4(bn2)*, a temperature sensitive mutant which limits germline development at 20°C or above, leading to sterility.

### Preparation of cohorts for imaging, survival and healthspan assays

For imaging and survival assays, animals were synchronised by bleaching gravid adults and placing eggs on NGM seeded with *E. coli* OP50, and incubated at 15°C. At L4 stage animals were transferred to experimental plates seeded with *P. aeruginosa* wildtype, *P. aeruginosa* reporter strains or *E. coli* OP50 control and shifted to 25°C. For healthspan assays, animals were synchronised by egg lay and transferred to experimental plates and 24°C at L4 stage. All experimental plates were seeded with 250 μL of bacterial culture and used the following day.

### Microscopy and imaging

Animals were imaged at 24 and 96 hours after shifting L4s to experimental plates and 25°C. At each time point, 10-15 animals were mounted onto 2.5% agar pads and anesthetized with 25 mM tetramisole. DIC and fluorescence (488nm Exc./505-575nm Em) images were acquired with a CellCam Rana 200CR camera and Leica DMR compound microscope driven by Micro-Manager Studio Version 2.0.0. Two biological replicates were performed.

### Killing assays

Animals were shifted to experimental plates and 25°C at L4 stage and scored dead or alive at 48, 96, and 192 h. Animals were considered dead if not responsive to prodding with a platinum wire. Three replicates were performed with 30 worms per replicate. Animals with internal hatching or that went missing from the plates were censored.

### *C. elegans* healthspan assays

Healthspan was assessed using the WormGazer™ automated imaging technology (Magnitude Biosciences). 60 mm plates were imaged using Raspberry Pi Version 2 cameras at a distance of 60 mm from the plate using white transmission illumination from a generic LED light panel. The cameras were located inside a temperature controlled laboratory set to 24°C. For each dish, a sequence of 200 images were taken over a 160 secs, with recording performed every 5 minutes until day 10 of adulthood. From these images, the number of moving objects is calculated by applying a threshold of the minimum speed of each object of 10 μm s^−1^. The speed is derived from the length of the object divided by the 160-s time interval of the imaging window. Plates were censored if they failed a quality control inspection after the experimental runtime, for example if they were contaminated with another microbe or the worms had burrowed into the agar. Censored plates were omitted from data analysis. Animals were imaged continuously using a minimum of 6 plates per condition.

## RESULTS

### *P. aeruginosa* biofilm formation *in vitro* is dependent on QS systems

We assessed the biofilm forming capacity of bacterial fluorescent reporters of the RhlI/R QS system and the virulence factor ExoS (Table 1). For RhlI/R we used a fluorescent transcriptional RhlA reporter fusion (*PrhlA-mNeonGreen*) in the PA14 background that targets expression during biofilm formation and infection. We also used the same reporter carrying a deletion in RhlR (*ΔrhlR PrhlA-mNeonGreen)*. The Δ*rhlR* mutant has been reported to have biofilm abnormal morphology phenotypes and attenuated virulence(Mukherjee et al., 2017). For ExoS, we used a transcriptional reporter fusion to the *exoS* promoter (*PexoS-gfp*) in PA14 and PA103 backgrounds. As controls we used *E. coli* BL21, which does not form biofilms(Wang et al., 2020) and the PA14 and PA103 wildtype strains. PA14 is a highly virulent clinical isolate and PA103 is a QS defective LasR mutant strain producing defective biofilms(Gambello and Iglewski, 1991; Haley et al., 2012; Le Berre et al., 2008).

We first tested biofilm formation *in vitro* using the peptidoglycan stain Crystal Violet, a classical staining method used to measure biofilms(Campo-Pérez et al., 2023). The bacterial strains were inoculated in microtiter plates. After 24 hours, biofilm formation was evaluated by removing non-adherent bacteria, staining the adherent cells using Crystal Violet, and measuring absorbance at 550 nm. PA14 showed a 4.2-fold increase in absorbance in comparison to the biofilm incompetent *E. coli* BL21, in agreement with good biofilm forming ability. In contrast, PA103 had similar staining levels as *E. coli* BL21, confirming a low biofilm forming capacity (Figure 2a). The *PexoS-gfp* and *PrhlA-NeonGreen* reporters in PA14 background showed absorbance levels similar to PA14 wildtype. *ΔrhlR PrhlA-NeonGreen* had a 1.7-fold lower absorbance compared to *PrhlA-NeonGreen*, and PA103 and PA103 *PexoS-gfp* had absorbance levels similar to *E. coli* BL21, suggesting compromised biofilm-forming abilities in these strains (Figure 2b-c). These findings confirm the biofilm forming capacity of PA14 and central role of the RhlI/R quorum sensing systems for the formation of *P. aeruginosa* biofilms.

**Figure 2.**
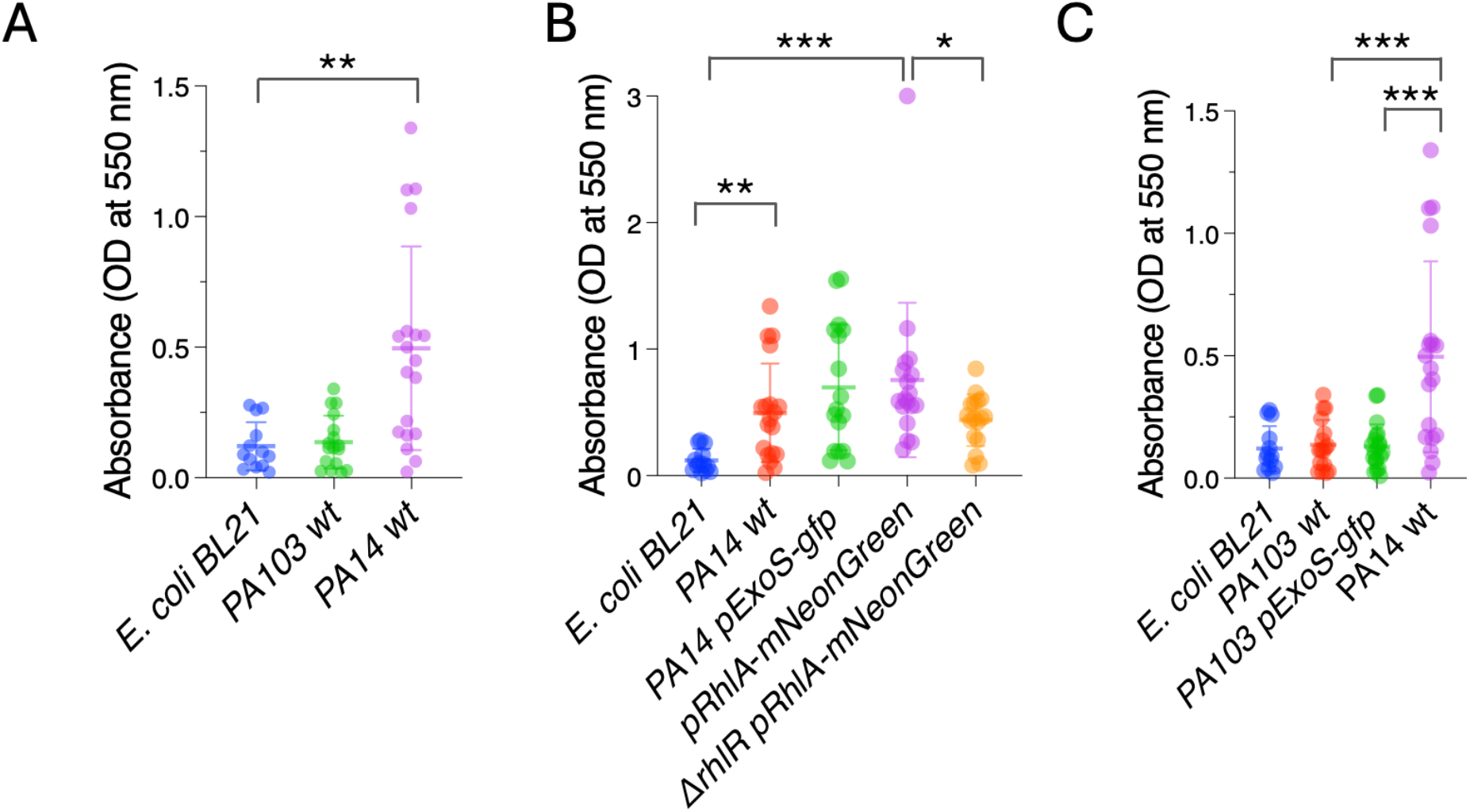
*P. aeruginosa* PA14 forms biofilms *in vitro*. Crystal violet staining was measured by absorbance at OD 550 nm in microtiter plates to quantify biofilm formation. A) PA14 has high biofilm-forming capacity compared to *E. coli* BL21. B) Reduced biofilm-forming capacity in *ΔrhlR PrhlA-NeonGreen*. C) Low biofilm-forming capacity in PA103 and PA103 *PexoS-gfp*. Three biological replicates per strain were performed. Error bars represent SD of three replicates. One-way ANOVA, *p <0.05, *p <0.01, **p <0.001, ****p <0.0001.

### *P. aeruginosa* reporters form fluorescent biofilms *in vitro*

Next, we tested if the biofilms can also be detected by measuring fluorescence produced by the reporters. Bacterial strains were inoculated in microtiter plates, and after 24 hours planktonic cells were washed off. Biofilms were quantified by measuring fluorescence at 488 nm/507 nm. *E. coli* BL21 and PA103 had low levels fluorescence relative to PA103 *PexoS-gfp*, which showed a 13.14-fold increase in fluorescence compared to PA103 background (Figure 3c). PA14 had significantly higher levels of fluorescence compared to *E. coli* BL21 (Figure 3b), likely due to autofluorescence resulting from pyocyanin production(Mukherjee et al., 2017). PA14 *PexoS-gfp* and *PrhlA-NeonGreen* fluorescence was 3.1 and 3.7-fold higher than *E. coli* BL21, respectively, but not significantly different from PA14 wildtype. *PrhlA-NeonGreen* fluorescence was reduced by 3.5-fold in the *ΔrhlR* mutant (Figure 3b).

**Figure 3.**
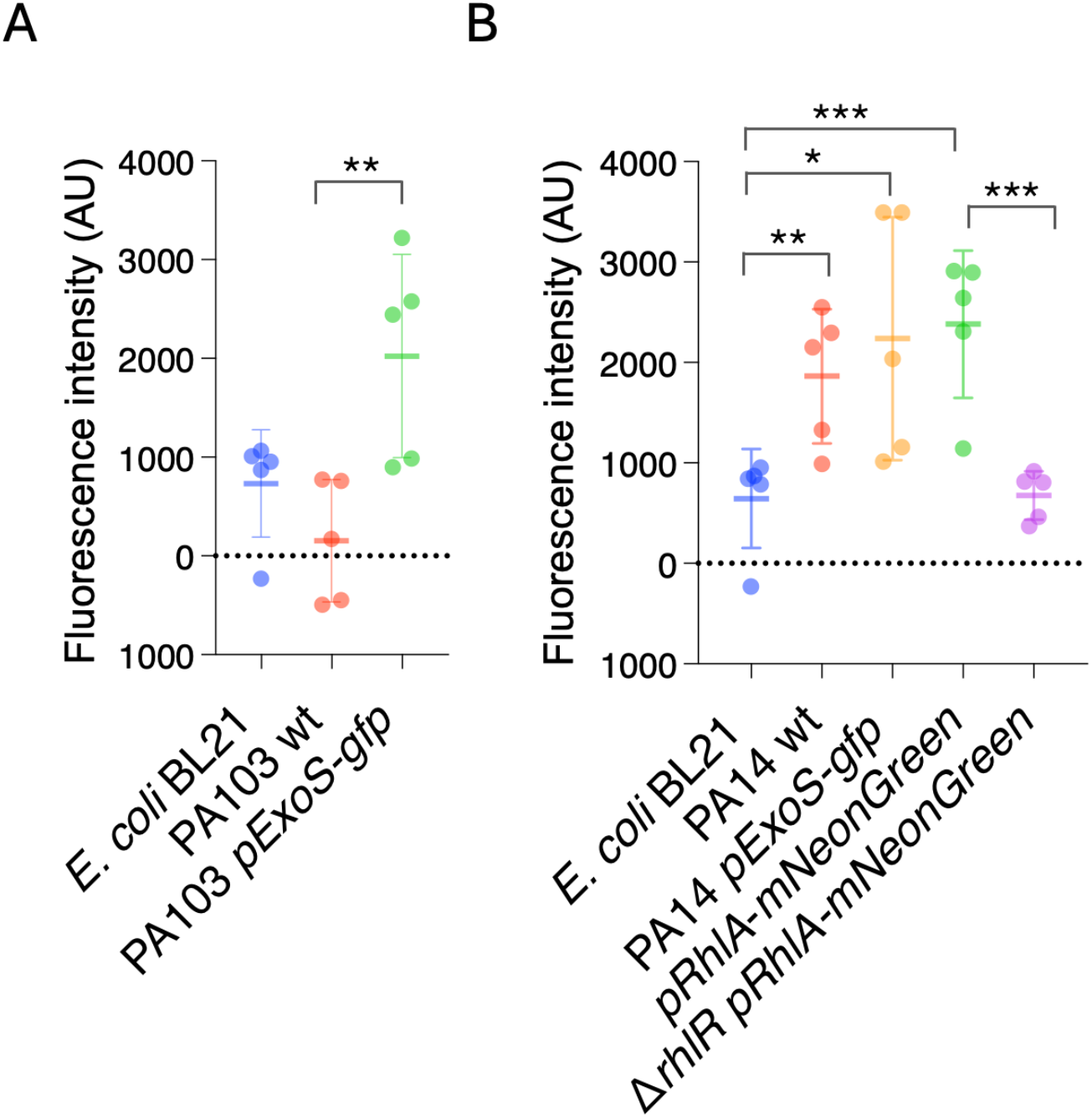
*P. aeruginosa* reporters form fluorescent biofilms *in vitro*. GFP fluorescence was measured in microtiter plates to quantify biofilm formation. A) Increased fluorescence in PA103 *PexoS-gfp* compared to PA103. B) Fluorescence in *PrhlA-NeonGreen* reduced by *rhlR* mutation. Three biological replicates per strain were performed. Error bars represent SD of three replicates. One-way ANOVA, *p <0.05, *p <0.01, **p <0.001, ****p <0.0001.

Except for PA103 *PexoS-gfp*, the reporters yielded similar results across both biofilm assays. The PA103 *PexoS-gfp* showed high levels of fluorescence compared to PA103, but similar levels of absorbance in the Crystal Violet assay, indicating discrepancies between the two methods for this strain. The fluorescence assay was not able to detect differences in between reporters and PA14 wildtype, likely due to overlapping spectra between the reporters and background fluorescence arising from autofluorescence.

### Infection by PA14 can be visualised using of ExoS and RhlA reporters

After measuring biofilms produced by the reporters *in vitro*, we asked if the reporters could be used to visualise bacterial infection *in vivo* in *C. elegans. C. elegans* is transparent, allowing imaging living animals without killing or dissection, making it ideal to monitor infection using fluorescent reporters. We cultivated *C. elegans* with the laboratory standard, non-pathogenic bacteria *E. coli* OP50 during development and exposed animals to the *P. aeruginosa* strains at L4 stage (end of development). Fluorescence was monitored *in vivo* using epifluorescence imaging 24 and 96 hours after exposure to *P. aeruginosa*. The later timepoint was chosen to capture early phases infection. We also attempted to image 6 days after exposure but found that these timepoints were not suitable, since for some of the conditions the surviving animals were very fragile, leading to bursting and death when mounted on slides.

In *C. elegans* the primary entry point for bacteria and site of infection is the intestine via the upper part of the gastro-intestinal tract, which consists of the mouth and pharynx. The intestine contains lysosome-related granules that generate autofluorescence. We observed fluorescence within the gut granules in all experimental conditions (Figure 4). In animals exposed to *E. coli* OP50, PA103 wildtype or PA14 wildtype, fluorescence was not seen in the intestinal lumen nor in other tissues. We did not observe major differences in fluorescence between the experimental conditions after 24 hours of exposure. In contrast, after 96 hours different patterns of fluorescence could be observed. In animals infected with PA14 *PexoS-gfp*, fluorescence could be seen in the intestinal lumen, consistent with it coming from GFP derived from bacteria proliferating in the lumen. In some animals widespread fluorescence, not only localised to gut granules, was present in the intestine as well as in the body cavity, indicating GFP-expressing bacteria had crossed the intestinal barrier and entered other tissues. Animals exposed to *PexoS-gfp* in PA103 background showed fluorescence contained to gut granules, with a subset of animals also having fluorescence present in the intestinal lumen. No fluorescence was observed in the body cavity, suggesting a reduced ability of PA103 to translocate through the intestine and infect the animal compared to PA14. Animals exposed to PA14 *pRhlA-NeonGreen* had fluorescence present in the intestinal lumen, intestinal cells and body cavity. This pattern was not observed in animals exposed to *ΔrhlR* PA14 *pRhlA-NeonGreen*. Together our imaging data show that by 96 hours of exposure, PA14 can cross the intestinal epithelial barrier to invade tissues, while for PA103 background the bacteria remain within the lumen. Translocation by PA14 can be visualised using the *PexoS-gfp* and *pRhlA-NeonGreen* reporters and is dependent on *RhlR*, confirming quorum sensing is important for PA14 pathogenicity in *C. elegans*.

**Figure 4.**
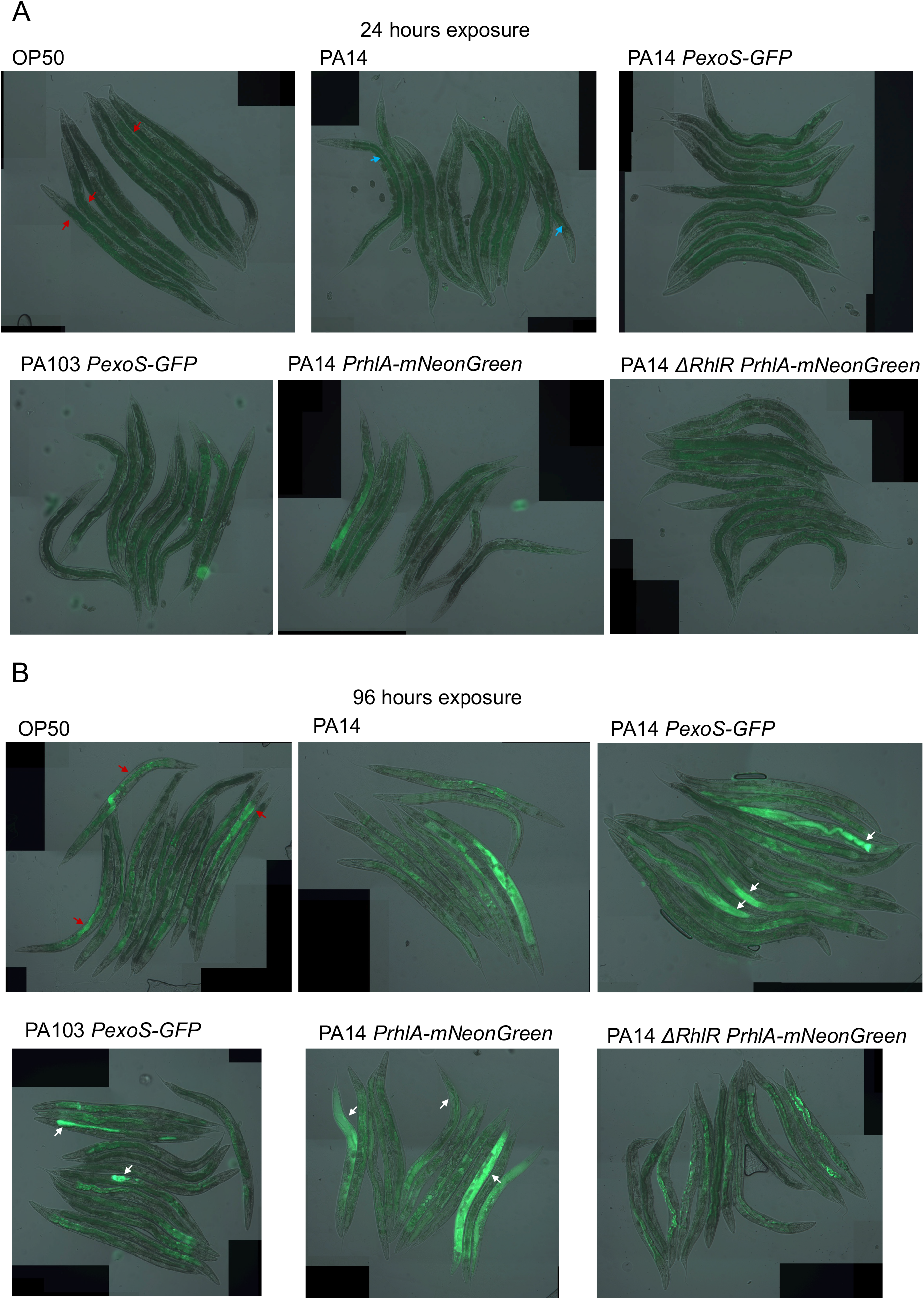
*In vivo* imaging of *C. elegans* exposed to *P. aeruginosa* reporter strains. **a**) Overlay of DIC and epifluorescence images taken after 24 hours of exposure. Autofluorescence from C. elegans gut granules within the intestinal cells (red arrows). Autofluorescence from *P. aeruginosa* pyocyanin within the intestinal lumen (blue arrows). **b**) Overlay of DIC and epifluorescence images taken after 96 hours of exposure. Widespread fluorescence observed in animals exposed to PA14 *PexoS-gfp* and *pRhlA-NeonGreen* (white arrows). Widespread fluorescence absent in *ΔrhlR* mutant background. PA103 *PexoS-gfp* fluorescence is contained within the intestinal lumen (white arrows). Autofluorescence from *C. elegans* gut granules within the intestinal cells (red arrows). Two biological trials were conducted.

### Reduction of RhlR signalling decreases PA14 pathogenic effects on lifespan and healthspan

We asked if the ability of the reporters to cross the intestinal barrier was reflected in the lethality of the strains. To determine if strains that translocate into the body cavity are more lethal than those that do not, we conducted killing assays. Animals were cultivated on *E. coli* OP50 and transferred to *P. aeruginosa* strains at L4 stage and survival was monitored at 48, 96 and 192 hours (8 days). Survival of control animals kept on *E. coli* OP50 was not affected during the experiment, and exposure to PA103 and PA103 *PexoS-gfp* resulted in a small, non-significant reduction in survival. In contrast, infection with wildtype PA14 and PA14 *PexoS-gfp* resulted in 73.2% and 71.6% of animals dead at 192 hours (Figure 5a). Exposure to PA14 *pRhlA-NeonGreen* resulted in 91.6% of animals dead while *ΔrhlR* reduced lethality of PA14 *PrhlA-mNeonGreen*, animals exposed to this strain had a survival of 61.4% at 196 hours (Figure 5b). These data are consistent with observations in our imaging experiments; PA103 is does not invade *C. elegans* tissues and has limited pathogenicity, while PA14 is capable of crossing the intestinal barrier and is highly pathogenic. PA14 pathogenicity is dependent on *RhlR*, consistent with RhlR signalling being important for virulence. Pathogenicity of PA14 *PexoS-gfp* and PA14 *PrhlA-mNeonGreen* were not different from PA14, thus expression of the reporters does not affect PA14 virulence.

**Figure 5.**
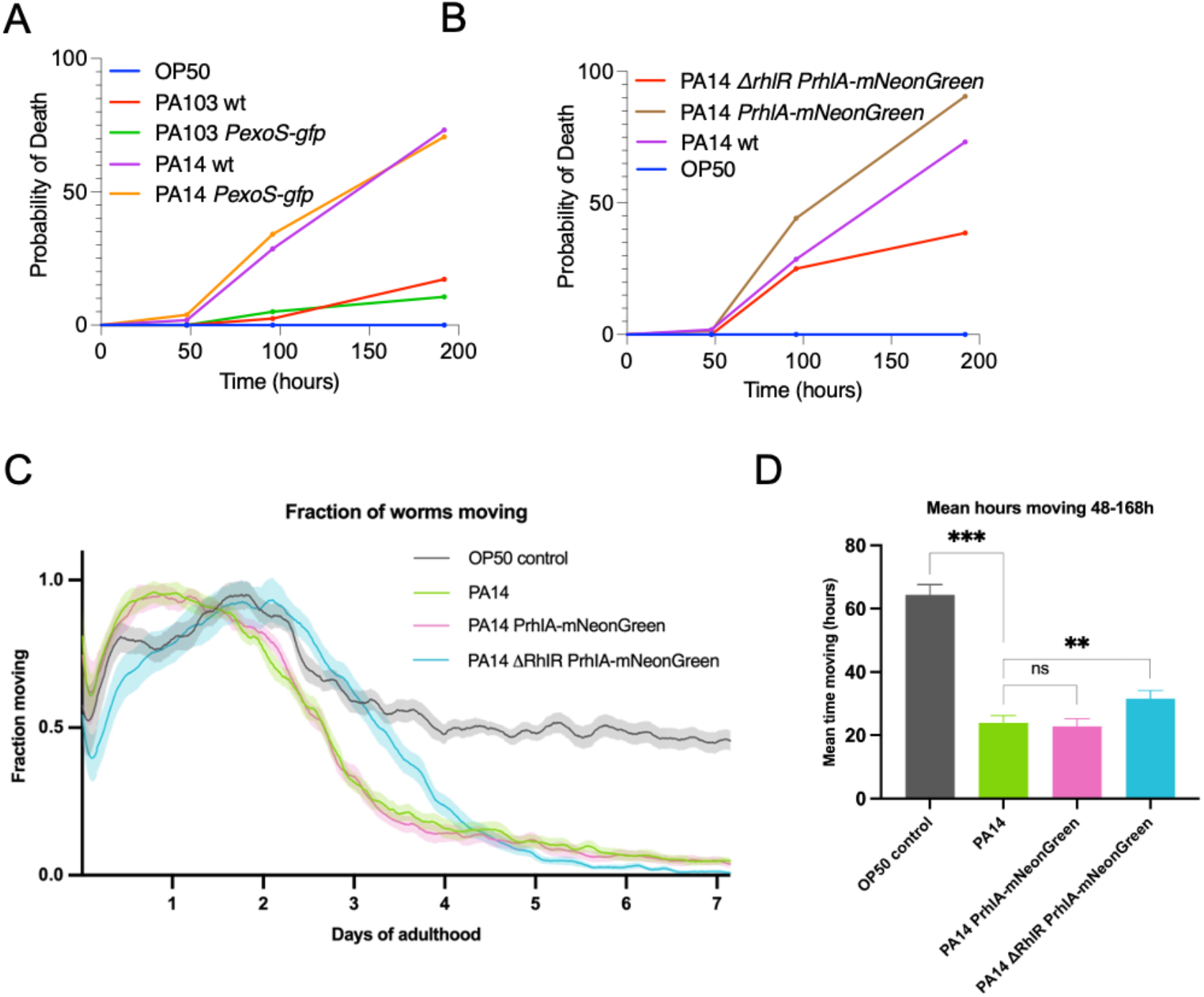
Decreased *RhlR* signalling in *P. aeruginosa* reduces pathogenicity in *C. elegans*. **a-b**) *C. elegans* survival was monitored up to 192 hours from L4 stage. Exposure to PA14 strains resulted in decreased survival, which was reduced by *ΔrhlR*. Exposure to PA103 did not affect survival. Three biological replicates with n=30 were performed. Log-rank (Mantel-Cox) test was performed for comparison of survival. **c-d**) *C. elegans* healthspan assays were performed by monitoring motility for 7 days starting at L4 stage. Exposure PA14 and PA14 *PrhlA-mNeonGreen* reduced motility compared to *E. coli* OP50. Animals exposed with to *ΔrhlR PrhlA-mNeonGreen* remained mobile for longer. **c**) Fraction of animals moving in healthspan assay. Shading shows SEM. **d**) The area under the curve integration for hours 48-168 of the healthspan assay. Error bars show SEM. Two biological replicates with n=180 were performed. **p<0.01, ***p<0.002, one-tailed t-test.

Infection models typically test effects on the ability of pathogens to kill the host, while effects on overall physical fitness are usually not assessed. We asked if PA14 pathogenicity in addition to affecting survival, also impacts healthspan. We used automated tracking and image analysis(Zavagno et al., 2024) to assess motility profiles of animals exposed to *P. aeruginosa* as a measure of healthspan. Animals exposed to PA14 wildtype, PA14 *PrhlA-mNeonGreen* and PA14 *ΔrhlR PrhlA-mNeonGreen* were tracked continuously from L4 stage until day 7 of adulthood to measure fraction animals moving, and hours spent moving and compared to *E. coli* OP50 controls. Animals in all conditions were active and mobile during the first two days of tracking, as expected in young, healthy animals (Figure 5c).

Animals grown on OP50 showed a partial reduction in movement between days 2 and 4 of adulthood due to normal age-related decline in muscle function(Huang et al., 2004; Newell Stamper et al., 2018). PA14 and PA14 *PrhlA-mNeonGreen* reduced overall movement and the time spent moving (Figure 5c,d). Animals exposed to PA14 *ΔrhlR PrhlA-mNeonGreen* exhibited a prolonged healthspan compared to PA14 and PA14 *PrhlA-mNeonGreen*, increasing both the distance moved and time spent moving (Figure 5c-e). Thus *P. aeruginosa* negatively affects both survival and healthspan and these effects are reduced when QS is compromised.

## DISCUSSION

*P. aeruginosa* biofilms have been studied extensively on glass or plastic surfaces *in vitro*, but much less is known about how biofilms form and develop on an epithelial barrier, the most common site of infection. In our study we infected the genetically tractable and transparent model organism *C. elegans* with fluorescent *P. aeruginosa* reporters to monitor infection *in vivo* while providing readouts of host health. By combining *in vivo* microscopy methods with new automated tracking technology, we show that the transcriptional RhlR receptor is important for the ability of *P. aeruginosa* to invade host tissues, and for pathogenic effects on survival and healthspan. As *P. aeruginosa* uses the same invasion mechanism across different host species, and mammals and invertebrates use conserved signal transduction pathways to activate defence-related genes(Tan et al., 1999), *C. elegans* is able to model certain aspects of mammalian pathogenesis. Our study is consistent with other studies examining *P. aeruginosa* infections and showing that QS plays a central role in virulence. For example, mutations in the RhlR transcription factor receptor alter biofilm morphology, reducing pathogenicity(Kumar et al., 2021; Mukherjee et al., 2018, 2017) and activation of host anti-pathogen defences(Peterson et al., 2023).

*In vivo* imaging infection of *C. elegans* with the *PrhlA-mNeonGreen* reporters showed that the widespread intestinal infection in wildtype background was absent in the animals exposed to the mutant, consistent with the compromised biofilm forming capacity in *rhlR* mutants resulting in reduced virulence. Using standard biofilm assays, we confirmed that the *rhlR* deletion reduces *P. aeruginosa* biofilms *in vitro. In vitro* the autofluorescence generated by pyocyanin production meant there were no differences in overall fluorescence levels between PA14 and the *PrhlA-mNeonGreen* reporter. In contrast, we could observe clear differences in pattern and distribution of bacterial fluorescence *in vivo*.

Also the *PexoS-gfp* reporter in PA14 background showed a high degree of biofilm formation capacity combined with pathogenicity. As with the *mNeonGreen* reporter, we could not detect differences in fluorescence intensity between wildtype PA14 and PA14 *PexoS-gfp in vitro* due to autofluorescence, but found major differences in bacterial fluorescence *in vivo*, with widespread tissue fluorescence following PA14 *PexoS-gfp* infection. This fluorescence was not observed in animals exposed to PA103 *PexoS-gfp*, and PA103 was also less pathogenic. In mice models of acute pneumonia, PA103 is highly virulent, severely reducing survival rates within 24 hours of exposure(Machado et al., 2010). The moderate pathogenicity of PA103 and lack of intestinal infection in our *C. elegans* assays highlight species-specific differences and raise the question of why *C. elegans* is able to avoid infection. A study using two different models of infection in *Drosophila* showed that PA103 results in high lethality in a fly nicking model but not in a feeding model(I et al., 2008). As PA103 lacks a functional QS system, QS might be required for lethality by feeding, e.g. to successfully attach to and infect intestinal cells. This can explain the lack of virulence in our *C. elegans* experiments were we also used feeding as means to expose the animals to the bacteria.

*P. aeruginosa* is listed as a high-priority pathogen in the 2024 WHO Bacterial Priority Pathogens List, due to its growing antibiotic resistance and global threat, especially in health-care settings(WHO Bacterial Priority Pathogens List, 2024). Currently very few treatments targeting *P. aeruginosa* biofilms are in development(Reynolds and Kollef, 2021) and the need for developing effective treatments against antibiotic resistant pathogens calls for *in vivo* models that are rapid and can be used in high throughput. Our work provides insight into how *C. elegans* combined with low-cost epifluorescence *in vivo* imaging and automated tracking technologies can be used to study *P. aeruginosa* infection. We envision the approaches described here being developed into high-throughput *in* vivo methods using e.g. 96-well plates and compound libraries, to screen for novel antimicrobials targeting *P. aeruginosa* biofilms, improving efficiency of pre-clinical antimicrobial testing and reduce costs, timelines and ethical burdens.

Future studies could make use of fluorescent reporters with emission wavelengths that do not overlap with those of pyocyanin and gut granules to more clearly differentiate between expression of the reporter and background. It would also be useful to develop methods that confirm the presence of biofilms in *C. elegans*, e.g. biofilm staining methods. Overall our work suggests that using *C. elegans* to study QS and biofilms could provide an exciting path forward for the development of novel antimicrobials.

## Author Contributions

Conceptualization M.E., D.W., R.H. and L.S.; methodology M.E. D.W., R.H. and C.S.; investigation S.A.B., M.R., F.X., M.F., S.M. and F.T.; resources M.E. and D.W.; data curation, M.R., F.X. and M.F.; writing—original draft preparation, M.E..; writing— review and editing, M.E., R.H. and D.W.; supervision, M.E., R.H. and D.W.; funding acquisition, M.E., D.W. AND L.S.; All authors have read and agreed to the published version of the manuscript.

## Acknowledgments

We thank Bonnie Bassler (Princeton University) and Gary Robinson (University of Kent) for bacterial strains used in the study. C. elegans strains were provided by the CGC, which is funded by NIH Office of Research Infrastructure Programs (P40 OD010440).”

